# Integrative multi-omics modelling for cultivated meat production, quality, and safety

**DOI:** 10.1101/2025.04.30.651459

**Authors:** Théo Mathieu, Sébastien Légaré, Astride Franks Nzekoue, Nathalie Jauré, Hannah Lester, Thomaz Dias, Remy Kusters

**Author notes:** Corresponding author *Email address:* (Remy Kusters). These authors contributed equally to this work (Théo Mathieu, Sébastien Légaré, Thomaz Dias, Remy Kusters).

## Abstract

Cell culture technology, which offers a promising solution for complementary food production, is slowly becoming a reality with the first wave of regulatory approvals in pioneering markets. However, significant challenges remain for large-scale cultivated meat commercialization, including effective scaling, cost efficiency, and product quality, but also scientific evidence to support regulatory approval and building consumer trust. In this paper, we discuss the potential of an integrative multi-omics approach to characterize and optimize cultivated meat production. By analyzing a network-based interactome model that integrates the transcriptomic, proteomic, and metabolomic layers, we achieve a system-level understanding of cellular metabolism and regulatory mechanisms. This approach can allow for precise monitoring and targeted interventions of critical quality and safety attributes associated with cellular biomass. We then describe a Target-Action-Metabolite (TAM) framework, which utilizes insights from the interactome to optimize cell culture conditions through actionable interventions. We illustrate the potential use of this framework through a case study involving Duck Embryonic Stem Cells (dESCs) for use in cell-cultured meat products, providing hypotheses for improving key metabolic pathways through targeted interventions on metabolites present in culture media. Finally, our paper highlights the potential of this interactome-based strategy to enhance bioprocess efficiency, improve product quality and ensure safety attributes, addressing regulatory challenges associated with cultivated meat production.

**Highlights:**

- Integrative multi-omics as a novel approach to support cultivated meat production.
- Interactome Models map molecular interactions to monitor and improve production.
- Multi-omics based strategy to scientifically support cell-culture safety assessment.
- Target-Action-Metabolites Framework to guide non-genetic interventions.
- Case study with avian interactome model to enhance cellular metabolism.

## 1. Introduction

Cultivated meat is rapidly developing to meet the growing worldwide demand for animal proteins (Da Silva & Conte-Junior, 2024). However, the path toward affordable and scalable cultivated meat still faces significant scientific and engineering challenges across the entire production pipeline, from cell line isolation to culture media formulation and bioprocess optimization (Zheng et al., 2025). Cultivated meat quality is defined by several key attributes, including nutritional composition, sensory properties (flavor, texture, color), and safety standards (Lambert et al., 2024). Unlike conventional meat, which derives its properties from the complex interactions between muscle structure, fat content, and post-mortem biochemical changes, cultivated meat must achieve comparable characteristics through carefully controlled cell culture conditions (Adi et al., 2024). Successful large scale commercialization of cultivated meat will not only require a cost-effective process but also robust scientific evidence supporting regulatory approval, safety and consumer acceptance for the end product (Chen et al., 2022). Integrative omics strategies, combining genomics, transcriptomics, proteomics, and metabolomics, offer a powerful framework for characterizing and validating cultivated meat at multiple biological levels (Lee et al., 2024). These approaches can help ensure the stability of cell lines, characterize cellular composition, identify potential allergens, metabolites and other bioactive compounds, and assess the bioavailability of key nutrients (Bakhsh et al., 2025). While many techniques can be adapted from pharma-oriented bioprocessing, cultivated meat production presents distinct challenges, including stricter cost constraints on culture media, constraints on the nutritional value of the cultured cells themselves, and a regulatory landscape that differs from that of pharmaceuticals (Gomez Romero & Boyle, 2023). Currently, integrative omics strategies are not widely used in novel food risk assessment, due to a lack of validated approaches although they represent a potential powerful tool to complement risk assessment and regulatory science. Moreover, the proactive use of such integrated omics, as advocated for in this paper, aligns with the paradigm of New Approach Methodologies (NAMs) that are encouraged for implementation by risk assessors such as the European Food Safety Authority (EFSA) (Junquera et al., 2024) to reduce animal testing and enhance product safety assessment (Westmoreland et al., 2022). In this paper, we explore how this multi-level systems biology approach can ensure a deeper understanding of the metabolic and regulatory state of the cultivated cells focussing on i) process development and media optimization, ii) quality and iii) safety attributes. In addition, we introduce a novel Target-Action-Metabolite (TAM) framework, providing a network-informed workflow for targeted interventions on the interactome, aiming to optimize cell culture conditions and media formulations, while monitoring potentially unwanted downstream impact on the interactome. By shifting from a classical gene-to-metabolism approach to a bi-directional metabolism-to-gene analysis, our framework has the potential to address some of the unique constraints of cultivated meat production, e.g. optimizing cellular pathways, through modifying downstream metabolites in the culture media rather than relying on genetic engineering which poses regulatory hurdles, and consumer acceptance issues, potentially hindering market adoption and scalability. We present some actionable hypotheses following from a case study on duck Embryonic Stem Cells (dESCs) used for cultivated meat production.

This paper presents a comprehensive exploration of our research, beginning with Section 2, which details the components of an interactome-based multi-omics model. Section 3 investigates the model’s potential in optimizing processes and media, ensuring quality, and assessing safety attributes. In Section 4, a case study on an avian cell line demonstrates how our custom avian interactome model reveals metabolic shifts during culture and how the TAM framework can safely enhance culture conditions and quality attributes. We conclude with a discussion on the future prospects of multi-omics modeling in cultivated meat production.

## 2. Integrative multi-omics

### 2.1. Benefits of interactomes

Our approach provides a framework for capturing the complex interactions between various molecular layers: genes, transcripts, proteins, and metabolites. By overlaying these layers, we gain a systems-level understanding of cell physiology, providing insights on how animal cells adapt and regulate their metabolism to culture conditions (e.g. physicochemical conditions, nutrient availability etc.). This allows us to identify, and act upon, the key regulatory genes/metabolites, protein-protein interactions and mechanisms that impact the growth kinetics and cellular identity over time. In the context of cultivated meat, we rely on a species-specific interactome model which allows us to highlight the biological transitions occurring throughout the whole production process and determine the molecular pathways responsible for these shifts. Figure 1 (A,B) shows a layout of the avian interactome used in this work. Details on the interactome construction are presented in Section 4.2.3.

**Figure 1:**
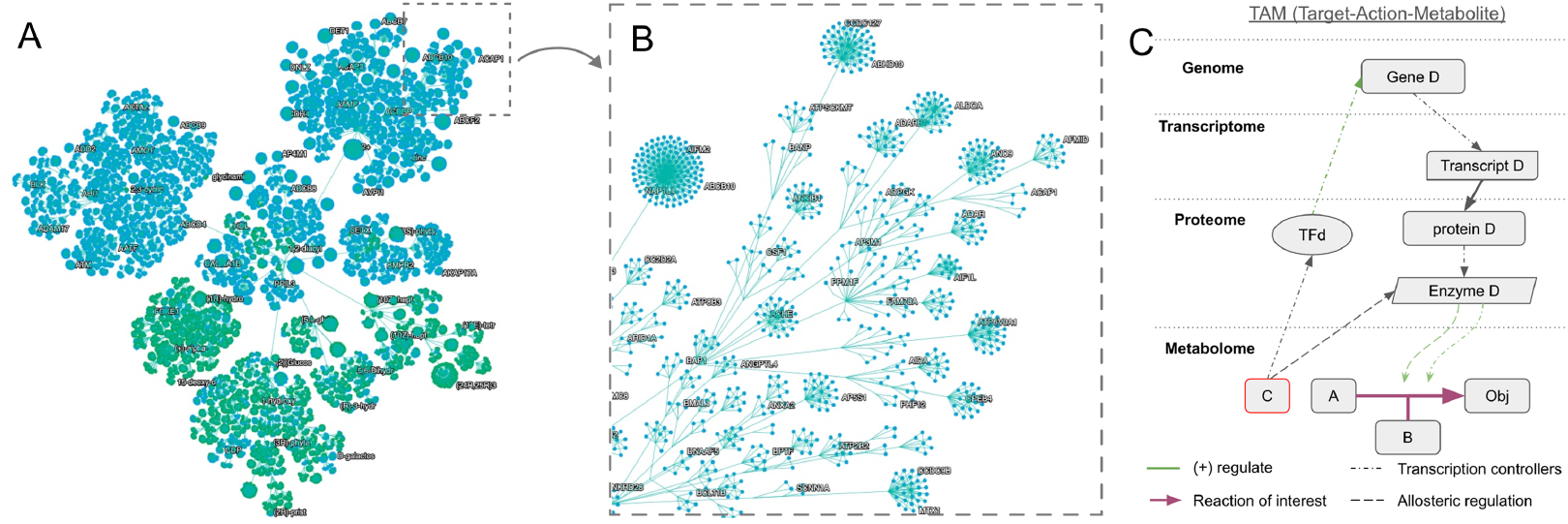
A) Layout of the avian interactome used in Section 4 using community detection hierarchical clustering to avoid the “hairball” visual. Gene/protein nodes appear in blue color and metabolite nodes in green. Labels show molecule identifiers for some nodes spread across the network. B) Zoom-in on the inset from A. Edges in this representation correspond to the hierarchical connections between clusters rather than molecular interactions. C) Two examples of bi-directional Target-Action-Metabolite: Transcriptional controllers and allosteric regulation.

### 2.2. Causal analysis

Causal analysis is used to uncover cause-and-effect relationships between molecular entities, such as genes, transcripts, proteins, and metabolites. Unlike traditional correlation-based methods, which identify associations without determining directionality, causal analysis aims to infer regulatory hierarchies and pinpoint key drivers of biological processes. In the context of the framework presented here, causal analysis allows dissecting regulatory hierarchies within the interactome, providing a causal directionality of action. While differential expression and pathway enrichment analyses provide valuable insights into cellular changes, they offer only a static view of molecular processes and do not reveal the full causal relationships between regulatory genes and metabolic adaptations. Causal inference methods help identify master regulators i.e. genes directly influencing metabolic shifts in avian cells, enabling targeted interventions for improving cell growth and metabolic efficiency. This approach offers three key advantages: (1) revealing connections to unmeasured molecules; (2) mapping interaction pathways from actionable molecules to otherwise unactionable differential elements; and (3) distilling extensive lists of differential genes/metabolites into manageable sets of actionable driver molecules. More concretely, for the case study in Section 4, this analysis enables the identification of transcriptional and post-translational regulators responsible for metabolic adaptation in the cell culture. This provides us with the opportunity to explore the full path across the interactome: metabolites, enriched pathways, prioritized differentially expressed genes (DEGs), and a set of key regulatory genes linked to metabolic adaptation through the TAM framework we present next.

### 2.3. Target-Action-Metabolite framework

The interactome model as described earlier serves as a powerful tool to characterize cellular function and its relation to cellular objectives, such as e.g. maximizing proliferation. Next we explore how we can perform directed actions on the interactome to e.g. improve the culture conditions. For this, we present the Target-Action-Metabolite (TAM) workflow which bidirectionally links Reaction (targets) Of Interest (ROI) to the Action Of Interest (AOI) through the various Precursors Of Interest (POI), directing the path across the interactome (Figure 1C):

- *Reaction (target) Of Interest (ROI)*: Define the cellular outcome to improve. This could be a specific metabolic reaction or pathway that correlates with a desired trait. For example, maximizing the rate of ATP production for faster cell proliferation, or increasing synthesis of a specific nutrient (like a polyunsaturated fatty acid) for nutritional quality. Generally these reactions are additionally constrained by substrate availability and thermodynamic feasibility.
- *Precursor(s) Of Interest (POI) identification*: Identify upstream factors in the interactome that strongly influence the target reaction. They can be considered the hidden variables in the interactome networks that functionally connect the ROI to the action of interest (AOI). Two examples are shown in Figure 1. Examples are i) Transcriptional controllers: TFs regulating ROI-associated genes and ii) Allosteric regulators, Metabolites modulating enzyme activity.
- *Actions Of Interest (AOI) determination*: Actionable interventions on the identified precursors. Actions could be genetic (e.g. overexpressing a gene), or environmental – such as adding a cofactor for an enzyme, altering pH or oxygen to shift metabolism, or supplementing a signaling molecule to trigger a pathway. In practice for cultivated meat, we favor actions that involve changing media components or process parameters (See examples in Figure 1).

Traditionally in pharma, AOI have focused directly on gene alteration, in order to impact the downstream metabolic layer, i.e., how gene expression influences metabolic processes and metabolite consumption. In cultivated meat production, regulatory frameworks and consumer concerns, particularly the conservative view on genetically modified food in the EU and the uncertainty of gaining approval, significantly restrict the direct genetic modification of cells. This hinders the pursuit of genetically modified approaches for cultivated meat, especially within the European market. This necessitates a broader approach that also considers how downstream metabolic conditions can affect upstream gene expression cascading further to overall cellular behavior (see Figure 1C for a schematic overview). By examining these bidirectional relationships across the interactome, we can pinpoint key metabolic shifts that drive cellular adaptation, thereby optimizing culture conditions more effectively.

## 3. Interactome models for cultivated meat

### 3.1. Meat quality attributes

Ensuring that cultivated meat products are safe, nutritious, and sensorially appealing is critical for consumer acceptance and market viability. Indeed, nutritional profiling is a key focus of novel food risk assessments to ensure that any new food product is not nutritionally disadvantageous. Moreover, quality attributes include not only safety and nutritional value but also sensory factors such as flavor, texture, aroma, and visual appearance, all of which are significantly influenced by the metabolites present in the cell culture media and their interactions within the cellular interactome. These metabolites can serve as precursors to flavor compounds and directly affect the final product’s sensorial profile. While to-date, studies looking at the link between cellular metabolites, proteome, pathway regulation and sensorial profiles are scarce, building larger scale interactome models will open the door to considerable innovation on this front. In conventional meat research, an integrative “foodomics” framework has emerged over the last decade. These multi-omics approaches (combining genomic, transcriptomics, metabolomics, proteomics, and volatilomics) provide deeper insights into the biochemical and molecular mechanisms underlying meat quality and enable the identification of specific biomarkers predictive of desirable traits such as tenderness, marbling, and flavor profile and intensity. Examples of such are e.g. the identification of predictive proteins biomarkers to understand and predict the sensory characteristics (tenderness, color, flavor) of meat products (Picard et al., 2017); the metabolomic characterization of beef reporting that the levels of acetyl-carnitine, adenine, beta-alanine, fumarate, glutamine, and valine were positively correlated with beef tenderness (Ramanathan et al., 2023); the identification and quantification of nucleotides (MP, GMP, AMP, inosine, and hypoxanthine) and acidic side-chain amino acids (Asp, Glu) as important contributors to the characteristic flavor profiles and umami taste of conventional meat (Hwang et al., 2020; Kawai, 2002). However, the previously presented interactome and network-based analyses allow for extending a similar foodomics approach even further by incorporating additional layers. In particular, the TAM framework presented, allows for the identification of specific AOIs relevant to the product’s quality attributes. By mapping complex interactions within cellular pathways, essential metabolites, gene expressions, and protein pathways can be identified, critical to meat quality attributes. Precise metabolic precursors can now effectively be targeted to impact these sometimes complex network attributes associated with particular quality attributes. The development of optimized cell culture media formulations, can now besides focussing on yield optimization, also consider additional metabolic and physiological pathways, leading to higher quality and sensorially appealing products. Examples worth pursuing could be adjusting lipid precursors in culture media, which for example, could shift cellular metabolism toward producing higher ratios of beneficial unsaturated fats, including omega-3 fatty acids (Post et al., 2020). Similarly, manipulating amino acid concentrations and specific metabolite levels can directly influence the sensory and nutritional outcomes of cultivated meats, providing control over texture, marbling, flavor formation, and nutritional profile (O’Neill et al., 2021).

### 3.2. Meat safety attributes

Among the potential food-safety hazards/risks related to cultivated meat and distinct from conventional meat production (Powell et al., 2025), the generation of novel secreted products with potential allergenicity and the occurrence of genetic instability or unintended cellular changes compared to the original cell type, are key considerations in regulatory risk assessment (Broucke et al., 2023). The interactome-based methodologies presented in the previous section enable systematic tracking of genome stability, expression profiles, and cellular characteristics across the entire production continuum, from cell isolation to final biomass. With increasing regulatory scrutiny and a growing demand for New Approach Methods (NAMs), data from these models can complement a comprehensive assessment of product safety. Indeed, EFSA envisions routinely using NAMs in risk assessments by 2030 to transition into a next-generation risk assessment that is exposure-led, hypothesis-driven, data-centric, and Adverse Outcome Pathway (AOP) based (Miccoli et al., 2022). Interactome models provide molecular fingerprints to verify cell line consistency and process control. For instance, analysis of the interactome can provide signatures that can inform cellular identity by comparing transcriptomic, proteomics, and metabolomic profiles across production batches, ensuring they remain within biological variability thresholds (Wang et al., 2023). Moreover, this temporal stability of cell multi-omic profile between successive batches, specifically at critical cell culture stages and during downstream operations, could offer a data-based approach to verify cell line stability within the cultivation process. As an example, in bovine satellite cell research, a developed genome-scale metabolic models have validated biomass predictions against experimental growth rates, demonstrating how multi-omics analyses can confirm the stability of metabolic pathways throughout muscle cell proliferation and differentiation processes (Lee et al., 2024). Similarly, network proximity metrics, such as kernel distance and separation distance, quantify interactions between cellular targets and critical pathways (e.g., glycolysis, Tricarboxylic Acid Cycle (TCA)), enabling predictive modeling of metabolic risks (Ma et al., 2024; Selevsek et al., 2020). Regulatory agencies responsible for the risk assessment of cultivated meat increasingly emphasize the importance of cell line documentation and culture media safety, advocating for harmonized standards in genetic stability assessments and media component categorization (APAC Regulatory Coordination Forum, 2024; Bakhsh et al., 2025). In a recent safety assessment, EFSA examined the safety of apple fruit cell culture biomass as a novel food and asked the applicant to provide a proteome profile analysis and comparison with conventional apple fruit. Proteome analysis of the Novel food revealed a marked difference between the proteins identified in the cell cultured biomass compared to conventional apples. EFSA noted that these differences could raise potential allergenic concerns, however, they still issued a positive safety conclusion. This highlights one of the first cases of the use of proteomic analysis in the risk assessment of novel foods. Network-based interactome models identify deviations in gene expression (e.g., toxin-related pathways) or metabolite accumulation, acting as early-warning systems for unintended cellular behavior (Todhunter et al., 2024). However, establishing universal “fingerprints” spanning all omics layers remains unachieved due to data heterogeneity and lack of validated (network-wide) biomarkers, requiring rigorous experimental verification (Lee et al., 2024). Pharmaceutical R&D provides precedents for multi-omics standardization: frameworks like MOGONET integrate multi-omics data to predict cellular subtypes, a strategy adaptable to cultivated meat for classifying cell line stability (Wang et al., 2023). Future advancements could adopt third-generation transcriptomics and 4D proteomics to enhance detection sensitivity, coupled with interactome-driven optimization to align cultivated meat profiles with conventional meat attributes (Todhunter et al., 2024). While multi-omics and network models promise robust safety assessment tools for regulators, enabling predictive risk analysis and batch-to-batch reproducibility, their full potential hinges on resolving data integration bottlenecks and establishing cross-industry validation protocols. Collaborative efforts will be critical to advancing these frameworks.

### 3.3. Process optimization & media development

Cell culture media optimization represents an important challenge in scaling cultivated meat production, requiring a rigorous understanding of metabolic and regulatory pathways. Indeed, with cell culture media representing a substantial production expense, optimization strategies are crucial to minimize these costs while improving production yield (Specht, 2020). The interactome model and the TAM framework we present here, allow to reveal and influence the complex interplay between cellular metabolism and environmental factors. By integrating these layers into predictive network models, we can identify key regulatory nodes and metabolic drivers that influence cell proliferation, differentiation, and product quality. To date, machine learning (ML) and systems biology are the emerging approaches for cell-culture media and bioprocessing optimization. ML, specifically Bayesian optimization, enabled efficient and cost-effective serum-free media design for C2C12 cell culture by intelligently exploring media component combinations to maximize cell growth with limited experimental data (Cosenza et al., 2023). In contrast, Genome-scale metabolic models (GEMs) provide a systems-level framework to simulate metabolic fluxes under varying media conditions. Building upon the success of bioproduction platforms like Chinese hamster ovary cell (CHO) cell cultures, which have demonstrated the efficacy of GEMs coupled with multi-omics data integration for improving biomass yield via media optimization, the cultivated meat sector is increasingly implementing systems biology approaches to develop customized cell line-specific media (Gomez Romero & Boyle, 2023). These GEMs reveal core pathways like glycolysis and TCA activity, enabling targeted media adjustments to enhance growth efficiency (Nikkhah et al., 2023). In a recent study, Lee et al., (Lee et al., 2024), demonstrated the feasibility of predicting and enhancing bovine cell culture processes via the application of a *Bos taurus*-specific GEM. By leveraging GEM and flux distribution analysis across varied media, this study examined glycolysis and the TCA cycle pathways in *B. taurus* cells, revealing active metabolic pathways for ATP and metabolite biosynthesis, while demonstrating the model’s ability to maintain essential metabolite biosynthesis through alternative routes when specific reactions are constrained. At the other side of the spectrum, Gene network inference (GNI) methods like dynGENIE3 (Huynh-Thu & Geurts, 2018) and Scribe (Qiu et al., 2020) decode regulatory interactions from multi-omics data, reducing high-dimensional gene expression matrices into interpretable network modules. In contrast to these studies, the approach we propose here, adopts a more holistic framework on the impact of media design. The POI which are often present in the metabolic layer, have direct upstream impacts on the proteomic and genomic layers through e.g., allosteric interactions. In the next section we will explore a concrete case study, covering the glycolysis pathway. Furthermore, focusing on metabolic drivers by modeling upstream regulatory influences, as previously described, can facilitate the identification of feed supplements that enhance productivity, demonstrating the utility of leveraging regulatory-metabolic interplay in the development of tailored media formulation (Herrgård et al., 2006).

## 4. Case study: Avian cell culture

### 4.1. Curation of the interactome model & metabolic layer integration

#### 4.1.1. Cell culture

To showcase the usage of the proposed framework we present a case study with an in-suspension duck Embryonic Stem Cells (dESCs). The experimental design for this study followed a batch-mode cell culture system, with samples collected at day 1, 4, and 7 of a shake flask batch culture. Cells were cultured in batch mode and did not receive additional nutrient feeding to simulate the various stages of cell growth: early stage exponential phase (day 1), late stage exponential phase (day 4) and nutrient starvation/apoptosis (day 7). Additionally, samples of fresh media were collected as controls. Cell counting and viability assessment were performed using a NORMA cell counter to track cell proliferation and density. The experiment was performed in triplicate (n=3), with concurrent collection of multiple data streams.

#### 4.1.2. RNA sequencing

For transcriptomic analysis, total RNA was extracted from cultured cells using the NucleoSpin RNA Mini Kit for RNA purification (Macherey-Nagel, Ref. 740955.5). RNA sequencing (RNA-seq) was performed on the Illumina platform using 150 bp paired-end reads. Quality control of raw reads was assessed with FastQC (Andrews, 2010), revealing consistently high base quality across all samples. Notably, 92.92% of bases exceeded Q30, indicating high-confidence sequencing data suitable for downstream analysis. Reads were aligned to the duck reference genome (*Anas platyrhynchos* (mallard), Pekin duck breed, NCBI ref: GCF_015476345.1_ZJU1.0 (Zhejiang University, 2020)) using STAR (v2.7.11b) (Dobin et al., 2013) under default parameters. The resulting alignments were used for subsequent gene expression quantification and differential expression analysis.

#### 4.1.3. Metabolomics

Intracellular metabolomics and spent media analysis (SMA) were carried out using LC-MS with both targeted and untargeted approaches to evaluate metabolite consumption and secretion patterns. Samples were processed for metabolite extraction using established protocols (Hernández Bort et al., 2014; Sellick et al., 2011). Briefly, after cells and culture medium separation, intracellular and spent medium metabolites were extracted via liquid nitrogen/cold methanol quenching, followed by protein precipitation and methanol/water extractions. Metabolomic analyses were performed using a Thermo Scientific Vanquish Horizon UHPLC coupled to a Thermo Scientific Orbitrap Exploris 480 mass spectrometer. Chromatographic separations used a Kinetex F5 column (150 × 2.1 mm, 2.6 µm) with a water/acetonitrile gradient containing 0.1% formic acid. Full Scan Data Dependent analyses (FSddMS2) in positive and negative modes were performed with a resolution of 120K in FS and 30K in ddMS2. Thermo Scientific TraceFinder and AcquireX/Compound Discoverer were used for targeted and untargeted analyses, respectively (Cooper & Yang, 2024).

#### 4.1.4. Interactome model

The custom avian interactome was built by agglomerating and harmonizing protein and gene interaction data from close to a hundred databases including BioGRID (Oughtred et al., 2021), Reactome (Milacic et al., 2024), and IntAct (Orchard et al., 2014). A metabolic layer was incorporated using curated metabolic pathway databases, where enzymes provide connection points into the gene-centric interactome. Orthology information about various avian species was used to unify this avian interactome. Only interactions that were deemed direct, involving a physical connection rather than being mediated by potentially unknown intermediary molecules, were conserved. Information about directionality and sign, whether the interaction target gets up or down-regulated, was also preserved when available. When possible the type of interaction, whether transcriptional, post-translational or metabolic, was also tracked. The resulting network is displayed on a hierarchical clustering layout at Figure 1(A,B). It contains approximately 12 000 gene/protein nodes, 4 000 metabolite nodes and 145 000 high confidence interactions. This is comparatively more interactions than alternative chicken interactome models available in literature (Konieczka et al., 2009). By combining high quality molecular interactions that are direct, with directionality, sign and type, this avian interactome is readily amenable to causal analysis.

### 4.2. Description of the analysis performed on the model

#### 4.2.1. Gene Ontology Enrichment

To track metabolic shifts, RNA-seq differential expression analysis was performed across three phases: i) early stage vs. late stage exponential phase, ii) late stage exponential vs. nutrient starvation phase. Gene ontology enrichment analysis identified terms associated with metabolic adaptation. In Figure 2 we showcase a selection of under/overexpressed ontology terms relevant for cellular characterization and identification. Notably, we directly observe metabolic adaptation between early stage and late stage exponential exemplified by a significant over-representation of cellular response to glucose starvation (GO:0042149), while biosynthetic associated ontology terms, such as steroid biosynthetic process (GO:0006694), are under-represented. Given that day 7 represents a departure from typical culture conditions due to nutrient starvation, we expect the cell culture to significantly regulate pathways associated with stress response when compared to the late stage exponential phase (day 4). As shown in Figure 2, we indeed observe the activation of different response pathways such as differentiation or apoptosis as suggested by e.g. an over-representation of positive regulation of neuron differentiation (GO:0045666) and the under-representation of DNA integrity checkpoint signaling (GO:0031570).

**Figure 2:**
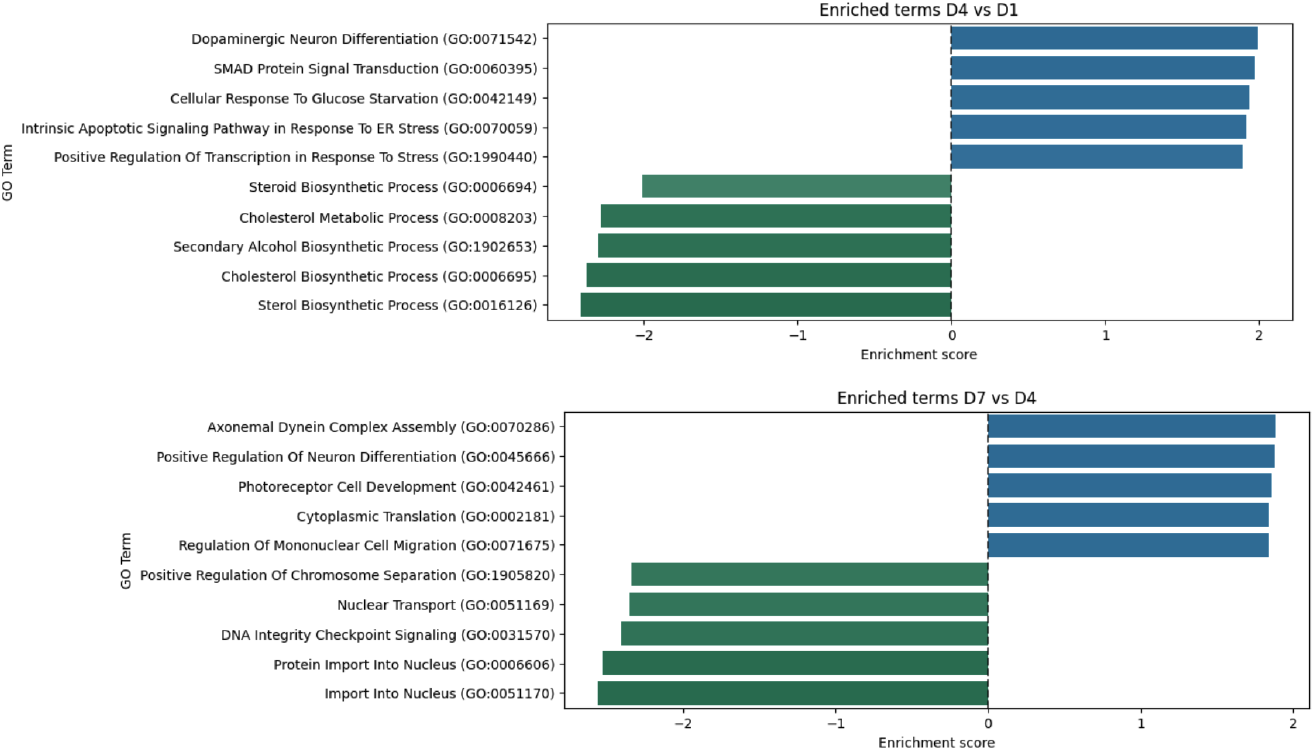
Main up and downregulated ontology terms at late-exponential stage versus early-exponential stage (top) and at nutrient starvation phase versus late-exponential stage (bottom)

#### 4.2.2. Focus on glucose and energy-related pathways

While cellular response to glucose starvation (GO:0042149) is enriched during the late-exponential stage compared to early-exponential stage, our metabolic data indicates that the substrate (glucose) is non-limiting (day 4 concentration > 25% of the initial concentration). This apparent contradiction prompted us to look more closely into glucose and energy-related pathways in the late-exponential stage compared to early-exponential stage. We found that the amount of intracellular glucose-6-phosphate as well as the expression of genes involved in glucose metabolism, like SLC2A1 and H2K, was increased in the late-exponential stage. At the same time, genes involved in the TCA cycle, like PDHA1, as well as lipogenesis genes like ACACA and FASN, were downregulated. This suggests some imbalance in cellular energy metabolic pathways, with an overconsumption of glucose and low oxidative phosphorylation.

#### 4.2.3. Causal analysis results

We then sought to find an AOI that could revert the aforementioned energy imbalance and potentially optimize cultivated meat production. We deployed causal analysis algorithms, leveraging the avian interactome and the TAM framework, to find genes and metabolites upstream of glycolysis, lipogenesis and the TCA cycle. Figure 3 displays the causal scores of transcriptional and post-translational regulators as violin plots. Figure 4 shows the relationships between significant causal genes implicated in the selected energy-related pathways. Gene MLXIPL, coding for the carbohydrate response element binding protein (ChREBP) is central to this causal network. At the top of the network are genes SIRT6 and PRKAA1, the catalytic subunit of AMPK, which are proposed as driver genes for observed differential transcriptomics and metabolomics data between the late-exponential and early-exponential stages.

**Figure 3:**
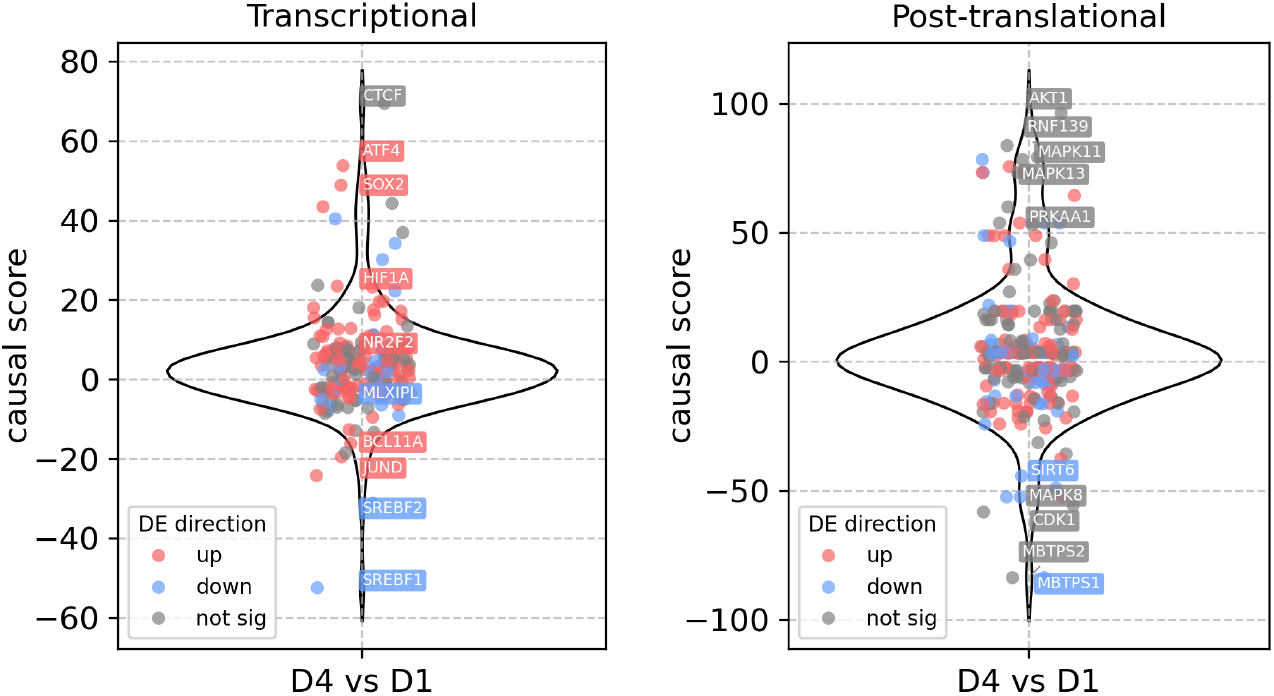
Violin plots of the transcriptional (left) and post-translational (right) causal analyses. Genes of interest to that study are labelled, including SIRT6, SREBF1, MLXIPL and HIF1A. Note that those genes do not necessarily have the highest causal scores, since we deliberately focus on energy metabolism and disregard genes involved in stress response or differentiation in the subsequent analysis.

**Figure 4:**
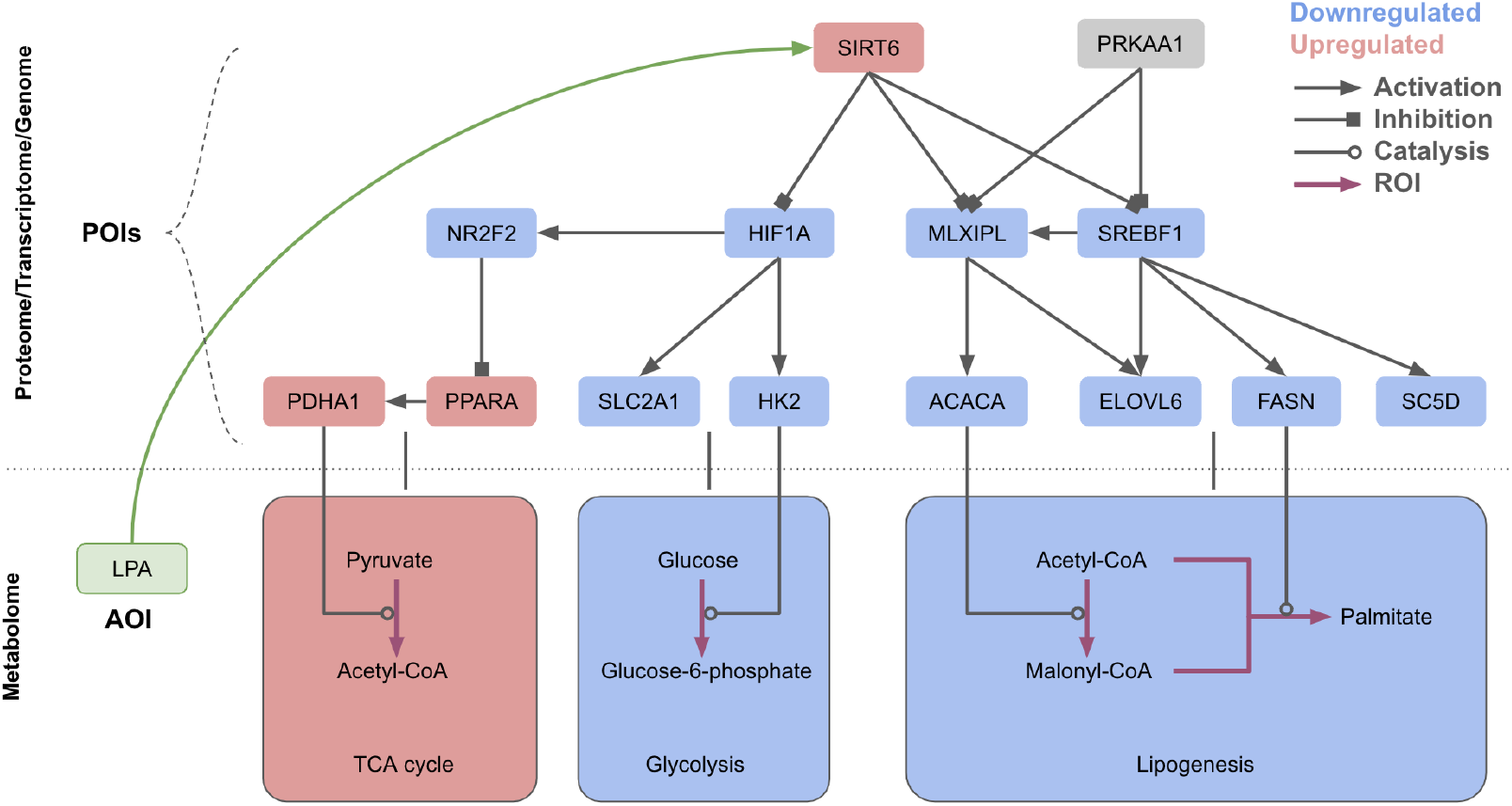
Causal analysis results represented as a hierarchical graph. The layout is split in two main layers with metabolites at the bottom and genes (implicitly standing for proteins and transcripts) at the top. Node colors do not correspond to the up and downregulation observed in the data, but rather to the expected effect induced by the AOI. Genes and biological processes expected to be upregulated are colored in red, downregulated in blue, and unaffected in gray. Proposed AOI metabolite LPA is displayed in green. Activation interactions are depicted as arrows-ended edges and inhibitions as square-ended edges.

#### 4.2.4. Proposed AOI

From the network of molecular interactions presented on Figure 4, it appears that activating SIRT6 could propagate downstream a signal to upregulate the TCA cycle and simultaneously downregulate glycolysis. As a side-effect, this could also decrease lipogenesis even more than the downregulation induced by PRKAA1, which may or may not be deleterious. SIRT6 can be activated by endogenous metabolites (Klein et al., 2020). This is particularly advantageous compared to typical knock-in or knock-out strategies (Wang et al., 2023) in the context of highly regulated cell cultures for food production. It allows us to propose a specific AOI; supplementing dESCs cultures with oleoyl-lysophosphatidic acid (LPA), or other metabolites highlighted in Klein et al. (2020). Supplementing LPA to culture media may activate SIRT6, shifting energy metabolism towards the TCA cycle, and possibly improving long term viability of the cultivated cells. This exemplifies how integrative systems biology and interactomes can be leveraged to uncover actionable targets to potentially optimize cultivated meat production.

## 5. Conclusions

In this paper, we advocate for the adoption of a multi-level systems biology approach, centered around avian interactome models and omics-driven analyses, to advance the characterization and improvement for cell lines used in cultivated meat production. By integrating genomic, transcriptomic, proteomic and metabolomic data, we gain a deeper understanding of cellular metabolism, regulatory mechanisms, and quality attributes. This approach will enable future interventions for optimizing cell growth, media development, and ensuring the safety and nutritional equivalence of cultivated meat. This proactive use of omics fits into the paradigm of New Approach Methodologies (NAMs) that regulators encourage to reduce animal testing and having a more holistic view of the changes occurring during the adaptation and production process. The case study of dESCs demonstrates the practical application of our framework, highlighting how integrative omics can identify key metabolic adaptations and regulatory genes. This approach not only enhances our understanding of cellular processes but also provides actionable insights for improving cell culture conditions and media formulations. The shift from a classical gene-to-metabolism to a bi-directional metabolism-to-gene analysis addresses some of the unique constraints of cultivated meat production by identifying nodes in the metabolic layer rather than acting directly on the genome through gene regulation. In the context of regulatory science, a recent review by Ong et al. (2023) highlights the importance of data sharing and standardized databases for foodomics would greatly benefit the field. There is currently a lack of comprehensive and wide-spead reference data on proteins and metabolites in the context of cultivated meat. Establishing databases for cultivated meat omics profiles, perhaps an open repository of genomic, proteomic, and metabolomic data from various companies and research labs, would help create benchmarks for cultivated meat at the molecular level. This can accelerate safety assessments by learning from public data on cell line performance. As regulatory frameworks for cultivated meat continue to evolve, integrating omics-based approaches into risk assessments and quality control processes will be crucial.

## 6. Declaration of artificial intelligence

During the preparation of this work the authors used Gemini in order to improve language. After using this tool, the authors reviewed and edited the content as needed and took full responsibility for the content of the publication.

## Acknowledgments

We thank Chloé Lopez, Louisa Barthélemy, João Palma and Meghiya Michelle for conducting the cell culture and sample preparation.

